# Multiple sources of Shh are critical for the generation and scaling of ventral spinal cord oligodendrocyte precursor populations

**DOI:** 10.1101/534750

**Authors:** Lev Starikov, Miruna Ghinia-Tegla, Andreas H. Kottmann

## Abstract

Graded Sonic Hedgehog (Shh) signaling emanating from notochord and floorplate patterns the early neural tube. Soon thereafter, Shh signaling strength within the ventricular zone becomes dis-contiguous and discontinuous along the ventral to dorsal axis suggesting a distribution of Shh that cannot be achieved by diffusion alone. Here we discover that sequential activation of Shh expression by ventricular zone derivatives is critical for counteracting a precocious exhaustion of the Olig2 precursor cell population of the pMN domain at the end of motor neuron genesis and during the subsequent phase of ventral oligodendrocyte precursor production. Selective expression of Shh by motor neurons of the lateral motor column at the beginning of oligodendrogenesis ensures a more yielding pMN domain at limb levels compared to thoracic levels. Thus, patterned expression of Shh by ventricular zone derivatives including earlier born neurons contributes to the scaling of the spinal cord along the anterior – posterior axis by regulating the activity of a select ventricular zone precursor domain at later stages of development.

## Introduction

The “Bauplan”, or body plan, of vertebrates can be recognized among members of the same species and across phyla despite significant differences in the absolute size of individuals of the same species or the form of different species. Chiefly responsible for our ability to recognize the common features of a body plan is the proportionate differentiation and growth of the constituent parts of an organism irrespective of absolute size. How developmental processes maintain a constant ratio of physical pattern features with changing size, a property known as scale invariance, is not completely understood (Huang and Umulis 2018).

Successful development is dependent on highly stereotyped series of inductive events that result in the determination of progressively increasing numbers of cell fates within rapidly growing tissues. Ensuring scale invariance could be achieved by mechanisms that are distinct from cell fate determination processes and could act at different developmental stages (Barkai and Ben-Zvi 2009, Umulis and Othmer 2013). Alternatively, patterning mechanisms themselves could be modified to generate a size-invariant output (Kicheva and Briscoe 2015). Cell fate determination is governed by a handful of cell signaling factors termed morphogens that are secreted from spatial fix-points in the developing embryo and form gradients of activity across a developmental field (Lander 2007). Morphogen signaling induces distinct transcriptional programs in naïve precursor cells dependent on signaling thresholds that are determined by signal strength and duration (Kicheva, Bollenbach et al. 2014). Experimental and theoretical studies revealed several mechanisms by which morphogen signaling can be scaled to embryo size (Gregor, Bialek et al. 2005, Houchmandzadeh, Wieschaus et al. 2005, Howard and ten Wolde 2005, McHale, Rappel et al. 2006, Ben-Zvi and Barkai 2010, Cheung, Miles et al. 2014, Uygur, Young et al. 2016). The commonality of these studies is that they define mechanisms of scale invariance across embryos during early stages of development. However, scaling occurs within the same embryo and throughout development as different body parts reveal proportionate changes in size regardless of when they are specified during development. How is proportionality ensured across developmental stages within the same embryo?

We have explored this question in the context of the switch from neurogenesis to gliogenesis in the developing neural tube. The spinal cord is a tube-like structure with enlargements at segmental levels that provide control of, and receive proprioceptive information from, limbs. The larger spinal cord at brachial (forelimb) and lumbar (hindlimb) levels compared to thoracic levels is chiefly the result of the presence of a greater number of motor neurons (MNs) that sub serve limb musculature, and correspondingly larger numbers of interneurons and glial cells (Bjugn and Gundersen 1993, Dasen 2017). Since the basic patterning and structure of the spinal cord is highly similar along the neuraxis, the difference in size between limb levels and thoracic segments allows comparative studies that might reveal mechanisms involved in scale invariance. MNs are among the very first neurons in the developing neural tube whose fate becomes specified while most interneurons and all types of glia develop later (Jessell 2000). Whether and how the earlier production of neurons plays a role in the proportionate differentiation of subsequent cell types is not well understood. MNs and oligodendrocyte precursor cells (OPCs) emerge subsequently from the same domain, the “pMN” domain, within the ventricular zone of the developing spinal tube (Bergles and Richardson 2015, Traiffort, Zakaria et al. 2016) suggesting that cell-autonomous as well as cell non-autonomous mechanisms could be involved in the sequential and proportionate differentiation of MNs and OPCs (Traiffort, Zakaria et al. 2016). The elucidation of the molecular control mechanisms that determine the precise “switch” from neurogenesis to gliogenesis of the pMN domain and the segment-specific yield of the pMN domain have been hampered by a scarcity of cell type selective- and temporally-specific genetic tools to dissect MN and OPC production within the pMN domain. Despite these difficulties, studies have implicated the morphogen Sonic Hedgehog (Shh) in regulating both, MN and OPC production (Briscoe and Ericson 1999, Traiffort, Zakaria et al. 2016).

Shh plays a critical role in MN and OPC differentiation at several developmental stages. First, graded Shh signaling originating from the notochord induces distinct transcriptional programs in overlying naïve neural ectoderm cells in a concentration-dependent manner that leads to the establishment of molecularly distinct precursor domains along the ventral to dorsal axis of the developing neural tube (Dessaud, McMahon et al. 2008, Alaynick, Jessell et al. 2011, Yu, McGlynn et al. 2013),. The transcription factors activated by Shh are responsible for determining the cell fates in the derivatives of these precursor domains. Thus, Olig2 expression marks MN precursors, (pMN) (Mizuguchi, Sugimori et al. 2001, Novitch, Chen et al. 2001), Nkx2.2 expression marks the more ventral V3 interneuron progenitors (p3), and Dbx1 is expressed in the pO domain located at the equator of the neural tube leading to interneurons situated dorsal to MNs (Briscoe, Pierani et al. 2000). Subsequently, persistent Shh signaling originating from the medial floorplate (MFP) located at the ventral midline of the developing neural tube is critical for maintaining the identities of each of these domains throughout neurogenesis (Dessaud, Ribes et al. 2010). The ventral neural tube expands rapidly during neurogenesis and Shh signaling strength measured by expression levels of Shh target genes declines progressively within the precursor domains (Balaskas, Ribeiro et al. 2012, Kicheva, Bollenbach et al. 2014). Surprisingly, the beginning of oligodendrogenesis is marked by increased Shh signaling strength within the pMN domain (Danesin, Agius et al. 2006). How this dynamic increase in Shh signaling is achieved subsequently to a decline in Shh signaling strength and whether it is of functional importance for OPC specification and production is an area of active research (Traiffort, Zakaria et al. 2016). Since the Shh gradient emanating from the MFP displays a constant decay length and does not scale with congruent and rapid growth during early development (Cohen, Kicheva et al. 2015), its influence on pMN domain activity must likely wane during neurogenesis. Accordingly, multiple adaptation mechanisms have been proposed to underlie temporally dynamic Shh signaling at the time of initiation of OPC production (Traiffort, Zakaria et al. 2016). One such mechanism is the accumulative storage of Shh in the extracellular matrix followed by Sulfatase 1 dependent release leading to a greater concentration of Shh than could be achieved by continuous production and diffusion (Danesin, Agius et al. 2006, Touahri, Escalas et al. 2012, Al Oustah, Danesin et al. 2014). While the genetic ablation of Sulfatase 1 curtails the production of OPCs, the current experiments cannot exclude the possibility that other signaling factors than Shh are released from the extracellular matrix by Sulfatase 1 and then participate in the regulation of OPC production. Another mechanism might be the sequential production of Shh by previously specified ventricular zone derivatives (VZD). Prominent sources of VZD_Shh_ are the lateral floorplate (LFP_Shh_) (Charrier, Lapointe et al. 2002, Park, Shin et al. 2004, Al Oustah, Danesin et al. 2014), which is constituted by cells emigrating from the p3 domain, and MNs (MN_Shh_) (Oppenheim, Homma et al. 1999, Akazawa, Tsuzuki et al. 2004). The selective ablation of Shh from LFP or MNs without impacting earlier patterning of the spinal cord has not been achieved and causal evidence for the involvement of LFP_Shh_ and MN_Shh_ in the regulation of OPC generation is therefore lacking.

Here we produced a series of mouse lines with different degrees and tissue selectivity of conditional Shh gene ablation from the MFP, LFP, and MNs during spinal cord development. We find that VZD_Shh_ is critical for the maintenance of the Olig2 expressing cell population in the pMN domain during the phase of oligodendrocyte precursor cell (OPC) production. While early neural tube patterning and MN development can proceed successfully in the absence of these Shh sources, the pMN domain exhausts of Olig2 expressing cells during neurogenesis leading to a subsequent and spinal level specific reduction in OPC production. LFP_Shh_ is needed for OPC production throughout the spinal neuraxis. In contrast, MN_Shh_, which occurs earlier at limb levels than thoracic levels, is critical for maintaining a larger pMN domain at limb levels compared to thoracic segments throughout OPC production. Our data provides causal evidence for the critical involvement of sequential and spinal level restricted expression of Shh by VZD for scaling of OPC production along the neuraxis.

## Results

### Shh expression in the ventral spinal cord

To examine the contributions of different sources of Shh in the ventral spinal cord to pMN domain activity at the time of the transition from MN to OPC production we first expanded on previous descriptions of Shh expression during spinal cord development. We used a gene expression tracer and conditional loss of function allele of Shh (Shh-nLZ^C/C^, abbreviated Shh^C/C^) in which a bicistronic mRNA is transcribed from the un-recombined Shh locus that encodes Shh and nuclear targeted LacZ (Gonzalez-Reyes, Verbitsky et al. 2012). The allele allows the quantification of the numbers of cells that express Shh and the determination of Shh ablation efficiencies in response to Cre activity with single-cell resolution. This approach reveals developmental stage- and spinal level-specific patterns of Shh expression and relative contributions to ventral Shh production from at least 4 distinct cell populations. At brachial levels at E12.5 we find Shh in all cells of the MFP (defined by co-expression with FoxA2 and situated at the ventral midline stacking 4-5 cells high along the ventral to dorsal axis), LFP (defined by reduced levels of FoxA2 and Shh compared to MFP, situated immediately dorsal to the MFP and stacking about 3-4 cells high along the ventral to dorsal axis), LFP* (defined by coexpression of Nkx2.2 and Shh and situated in part at the lateral edges of the LFP and in part as isolated cells flanking the p3 domain), and in 30% of all motor neurons (MNs) of the lateral motor column (LMC, defined by co-expression with Hb9 and lateral position in the ventral horns) (**Fig. 1**). In contrast, at thoracic and lumbar levels at E12.5, we find Shh expression only in MFP, LFP*, and LFP (**Fig. 1**). Based on numbers of nLacZ+ cells present at E12.5 we estimate that about 57% of Shh production occurs by MFP, 30% by the collective LFP and 13% by MNs at brachial levels while about 50% of Shh production occurs each by MFP and LFP at thoracic and lumbar levels (**Fig. 1B**). The onset of Shh expression in these tissues is sequential and overlaps with both MN and OPC production (**Fig. 1C**), suggesting the possibility to dissect each source’s Shh contribution to pMN activity.

**Fig. 1.**
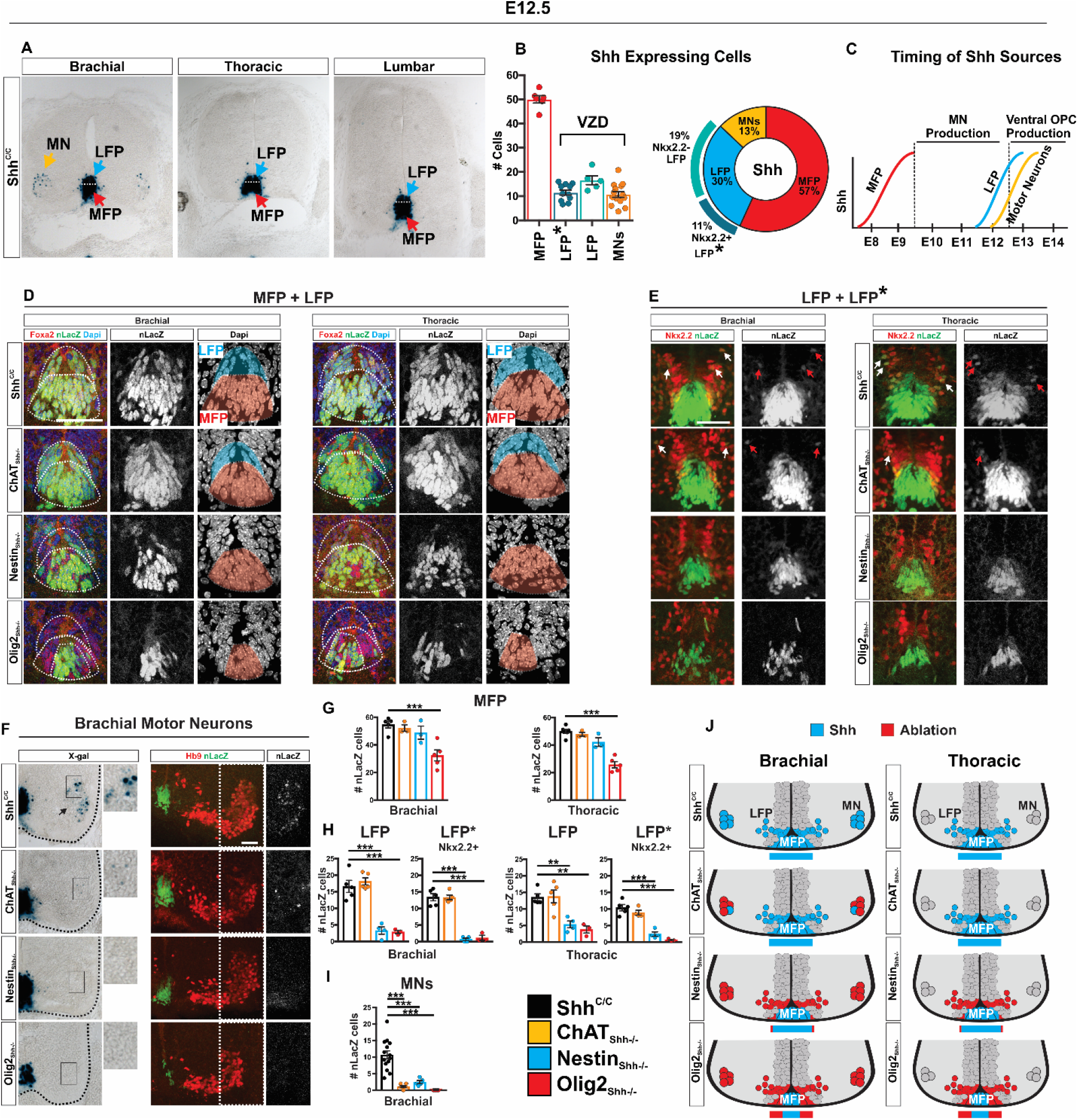
Shh expression and ablation strategy in the ventral spinal cord at E12.5. (**A**) X-gal staining of Shh expressing cells of E12.5 control Shh^C/C^ spinal cord sections. Three sources identified: Medial floor plate (MFP), lateral floor plate (LFP), and motor neurons (MN). (**B**) Numbers of nLacZ expressing cells in MFP (FoxA2+) n=6, LFP (FoxA2+ Nkx2.2-) n=5, LFP* (Nkx2.2+) n=11, and among MNs (Hb9+) n=14. Ventricular zone derived (VDZ): LFP, LFP*, and MNs. Breakdown percentages of each Shh source relative to total. Means ± SEM are shown. (**C**) Overlap in MN and ventral OPC generation from pMN domain with the timing of Shh expression from all sources. (**D**) Immunostaining for nLacZ and FoxA2 on brachial and thoracic segments. Identification of MFP (FoxA2+ nLacZ+) and LFP (FoxA2+ nLacZ+). LFP cells are identified as dorsal to MFP cells, oriented in tangent to MFP, and expressing lower levels of nLacZ. (**E**) Immunostaining for nLacZ and Nkx2.2 on brachial and thoracic segments. Identification of LFP (Nkx2.2-nLacZ+) and LFP* (Nkx2.2+ nLacZ+). Arrows point to migrating LFP* Nkx2.2+ nLacZ+ cells. (**F**) X-gal staining and immunostaining for nLacZ and Hb9 on brachial segments. (**G**) Quantification of MFP nLacZ recombination at brachial and thoracic segments for each Shh source per genotype. Brachial and thoracic segments Shh^C/C^ n=5, ChAT_Shh_^-/-^ n=3, Nestin_Shh_^-/-^ n=3, Olig2_Shh_^-/-^ n=5. Means ± SEM are shown. One-way ANOVA, Dunnett’s multiple comparison post hoc test. *p<0.05, **p<0.01, ***p<0.001. (**H**) Quantification of LFP and LFP* nLacZ recombination at brachial and thoracic segments for each Shh source per genotype. Brachial and thoracic segments Shh^C/C^ n=5, ChAT_Shh_^-/-^ n=4-5, Nestin_Shh_^-/-^ n=4, Olig2_Shh_^-/-^ n=3. Means ± SEM are shown. One-way ANOVA, Dunnett’s multiple comparison post hoc test. *p<0.05, **p<0.01, *** p<0.001. (**I**) Quantification of nLacZ recombination in brachial MNs. Shh^C/C^ n=14, ChAT_Shh_^-/-^ n=8, Nestin_Shh_^-/-^ n=5, Olig2_Shh_^-/-^ n=5. Means ± SEM are shown. One-way ANOVA, Dunnett’s multiple comparison post hoc test. *p<0.05, **p<0.01, ***p<0.001. (**J**) Schematic of Shh expressing cells at E12.5 brachial and thoracic segments and strategy of Cre ablation for each genotype. Scale bars, 50 μm.

At E13.5 lumbar LMC MNs begin to express Shh resulting in a pattern of expression that is qualitatively similar to brachial levels at E12.5 (**Fig. S2A**). By E14.5 medial motor column (MMC) MNs at all spinal segments begin to express Shh (**Fig. S1A**). At P20 Shh expression occurs in MNs, V0 cholinergic neurons, and remaining FP cells at all spinal cord segments (**Fig. S1B and S2C**). We observe a similar temporal and segmental pattern of Shh expression in the developing chick spinal cord (**Fig. S1C**).

To investigate the role of Shh signaling from these Shh sources onto the pMN domain, we generated a series of mouse lines with conditional and in part overlapping ablation of Shh. We used ChAT-Cre (ChAT_Shh_^-/-^) to target MNs, Nestin-Cre (Nestin_Shh_^-/-^) to target all Shh expressing VZD (MNs, LFP, and LFP*), and Olig2-Cre (Olig2_Shh_^-/-^), to target MNs, LFP, and MFP.

At E12.5 we did not observe Shh recombination in MFP cells in ChAT_Shh_^-/-^ mutants. Nestin_Shh_^-/-^ mutants had a few MFP cells which displayed Cre activity, but this ablation was insignificant (**Fig. 1D and 1G**). However, in Olig2_Shh_^-/-^ mutants the Shh recombination efficiency among MFP cells in brachial and thoracic spinal segments revealed a significant ~44% and ~39% loss resp. (**Fig. 1D and 1G**). We found no recombination of Shh in the LFP or LFP* in ChAT_Shh_^-/-^ at brachial or thoracic segments (**Fig. 1D, E, H**). Consistent with the previous reported expression of Nestin-Cre in all VZD between E10.5 to E12.5 (Kramer et al., 2006), and transient expression of Olig2-Cre in all VZD of the pMN domain and ventral to it (Ribes and Briscoe, 2009), we find near complete (~80%, and ~90% resp.) ablation of Shh from the LFP and LFP* in both Nestin_Shh_^-/-^ and Olig2_Shh_^-/-^ in brachial segments (**Fig. 1D, E, H**). At thoracic segments LFP recombination was less efficient at ~60% and ~72% for Nestin_Shh_^-/-^ and Olig2_Shh_^-/-^ resp. LFP* recombination was ~77% and ~94% for Nestin_Shh_^-/-^ and Olig2_Shh_^-/-^ resp. (**Fig. 1D, E, H**). Cre efficiency in MNs was about 80% in ChAT_Shh_^-/-^ and Nestin_Shh_^-/-^ mutants, and 98% for Olig2_Shh_^-/-^ (**Fig. 1F and 1G**). The location of Shh expression and the varied degrees of ablation of Shh at E12.5 are schematically summarized in **Fig. 1J**.

Using a conditional reporter allele (**Fig. 2A**), we find that Olig2-Cre but not Nestin-Cre is active in the MFP prior to notochord (NC) regression at E10.5 resulting in the ablation of Shh from 46 ± 5.1% and 8.5± 2.8% resp. of FoxA2+ cells (**Fig. 2B and 2C**). These results reveal that the degree of ablation of Shh from MFP in Nestin_Shh_^-/-^ and Olig2_Shh_^-/-^ animals is established at the time of MN production and remains fixed. The drastic loss of Shh expression in the MFP already at E10.5 in Olig2_Shh_^-/-^ animals prompted us to ascertain a possible patterning defect along the ventral midline. Consistent with previous reports that Shh expression by the notochord (NC) is sufficient for the establishment of precursor domains in the ventral spinal cord (Dessaud, Ribes et al. 2010, Yu, McGlynn et al. 2013) we find that the relative location and size of the p3- (Nkx2.2), pMN-(Olig2) and pO-(Dbx1) domains, and the location of the ventral border of the Pax6 expression domain are indistinguishable between Olig2_Shh_^-/-^ and control at E10.5 (**Fig. 2D and 2E**).

**Fig. 2.**
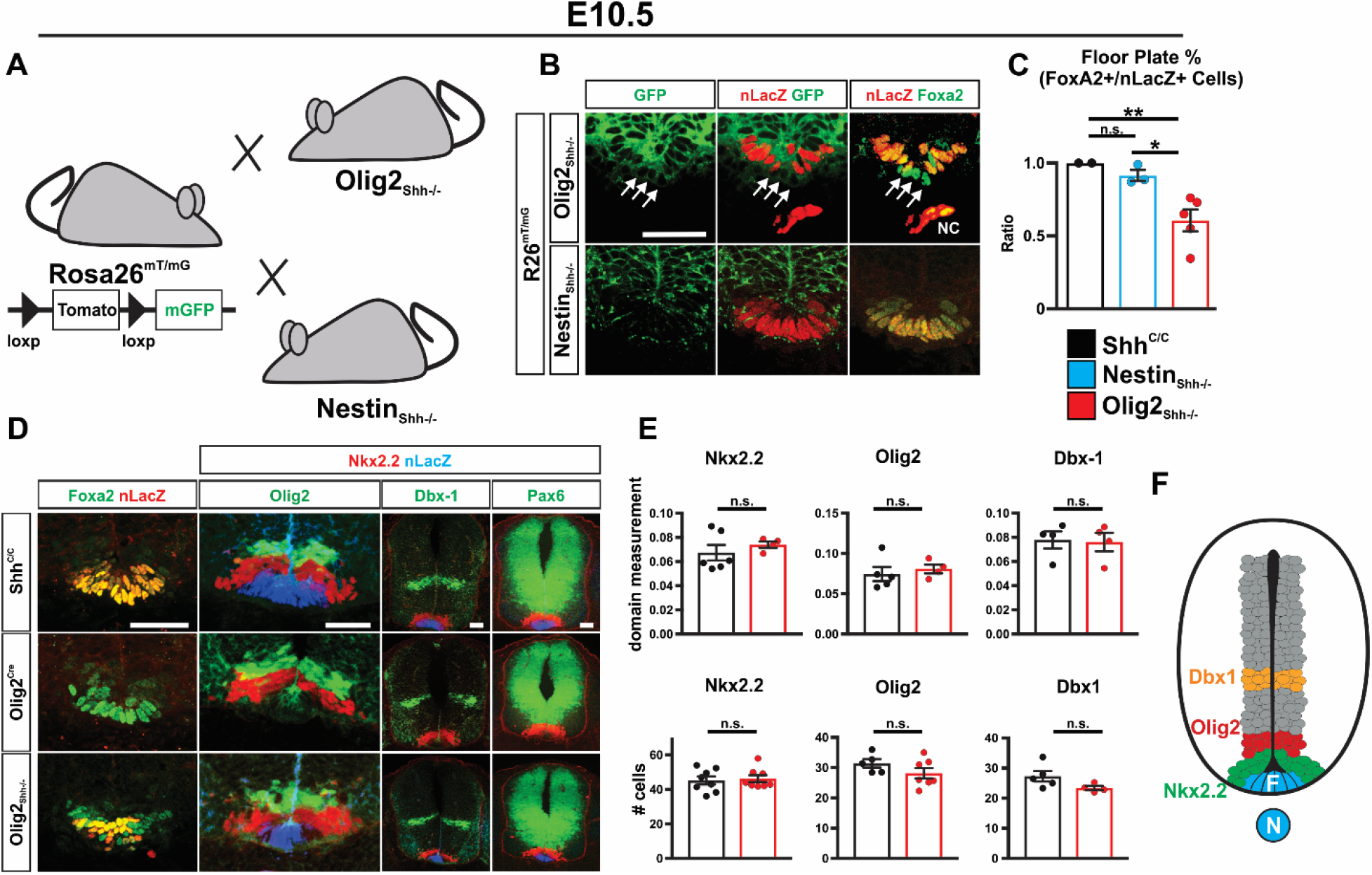
Early spinal cord patterning is unaffected in Olig2_Shh_^-/-^. (**A**) Detection of Cre activity in Olig2_Shh_^-/-^ and Nestin_Shh_^-/-^ mutants at E10.5 using the conditional R26mT/mG allele. (**B**) GFP co-labeling of Cre activity in addition to a loss of nLacZ from Foxa2+ cells demonstrates Shh ablation in MFP of Olig2_Shh_^-/-^ but not Nestin_Shh_^-/-^ embryos. Arrows indicate MFP FoxA2+ cells that have lost nLacZ expression. NC, notochord. (**C**) Quantification of recombination frequency of the ratio of Foxa2+ nLacZ+ double positive cells. Shh^C/C^ n=2, Nestin_Shh_^-/-^ n=3, Olig2_Shh_^-/-^ n=5. Means ± SEM are shown. One-way ANOVA, Tukey post hoc test. *p<0.05, **p<0.01. (**D**) Immunostaining of shh-sensitive domains, Olig2, Nkx2.2, Dbx1, and Pax6 are unaffected at E10.5 in Olig2_Shh_^-/-^, despite MFP recombination. (**E**) Quantification of relative domain sizes and numbers of Nkx2.2, Olig2, Dbx1. Domain measurement Shh^C/C^ n=4-6, Olig2_Shh_^-/-^ n=4. Means ± sEM are shown. Cell counts Shh^C/C^ n=5-8, Olig2_Shh_^-/-^ n=4-8. Data were analyzed by Student’s t test. *p<0.05, **p<0.01. (**F**) Scheme highlighting position of p3 (Nkx2.2), p0 (Dbx-1), and pMN (Olig2) domains relative to the MFP at E10.5.

Together, the varied degrees of quantifiable and source selective ablation of Shh from the ventral spinal cord and the preservation of early ventral tube patterning indicated that this set of recombinant mouse lines might be informative in the investigation of the effect of VZD sources of Shh onto the pMN domain during MN and OPC generation.

### Shh signaling from MFP, but not VZD sources influences MN generation

We investigated whether ablating Shh from VZD sources would impact the generation of MNs from Olig2 precursors of the pMN domain. We analyzed columnar pattern, relative distribution of MNs among columns, and absolute numbers of MNs of different columnar identity at brachial and thoracic levels. We first visualized MN columnar organization by immunostainings for MMC (Hb9+ Lhx3+), LMC_M_ (Hb9+ Isl1/2+), and LMC_L_ (Hb9+, Isl1/2-, Lhx3-) at brachial (**Fig. 3A**) and MMC (Hb9+, Isl1-), HMC (Hb9+, Isl1/2+), and PGC (Hb9-, nNos+) at thoracic (**Fig. 3E**) and found no apparent differences in the staining pattern either among ChAT_Shh_^-/-^, Nestin_Shh_^-/-^, Olig2_Shh_^+/-^, or Olig2_Shh_^-/-^ compared to Shh^C/C^ controls. Further supporting unaffected MN positioning and patterning, we find inconspicuous ventral root formation in Olig2_Shh_^-/-^ compared to Shh^C/C^ controls at E10.5 and E12.5. (**Fig. S3**).

**Fig. 3.**
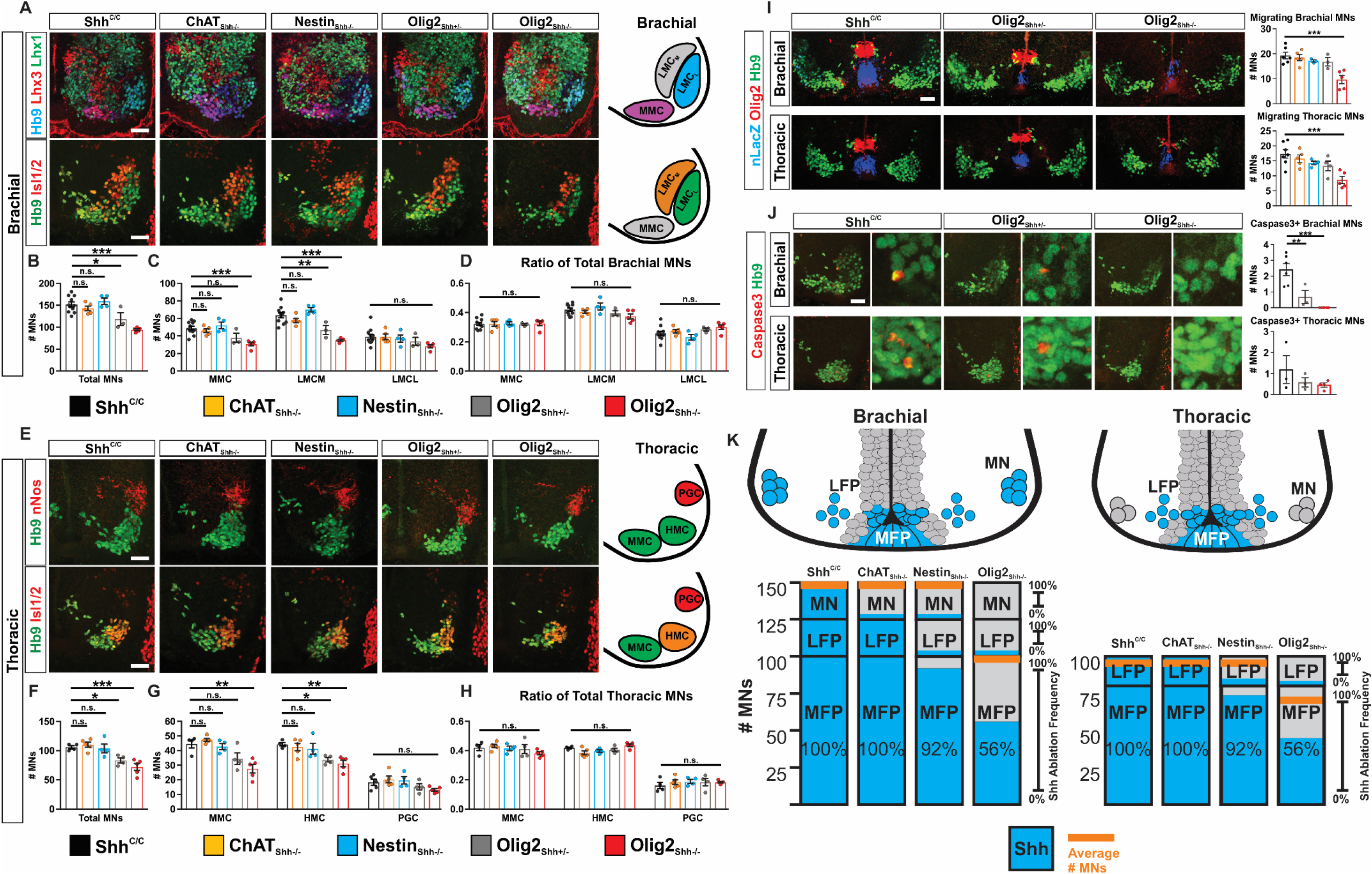
Shh signaling from MFP, but not VZD influences MN generation. **(A)** E12.5 brachial sections immunostained with Hb9, Lhx3, and Lhx1 to distinguish MMC and LMCL columns, and Hb9 and Isl1/2 to distinguish LMCM and LMCL columns. (**B**) Quantification of total brachial MNs. Shh^C/C^ n=11, ChAT_Shh_^-/-^ n=5, Nestin_Shh_^-/-^ n=4, Olig2_Shh_^+/-^ n=3, Olig2_Shh_^-/-^ n=5. Means ± SEM are shown. One-way ANOVA, Dunnett’s multiple comparison post hoc test. *p<0.05, **p<0.01, ***p<0.001. (**C**) Quantification of total numbers of MMC, LMCM, and LMCL MNs. Means ± SEM are shown. One-way ANOVA, Dunnett’s multiple comparison post hoc test. *p<0.05, ** p<0.01, ***p<0.001. (**D**) Ratio of each motor column to total brachial MNs. Means ± SEM are shown. One-way ANOVA, Dunnett’s multiple comparison post hoc test. NS, not significant, P>0.5. (**E**) E12.5 thoracic sections immunostained with Hb9 and nNos to distinguish MMC and HMC from PGC column, and Hb9 and Isl1/2 to distinguish MMC and HMC columns. (**F**) Quantification of total thoracic MNs. Shh^C/C^ n=4-5, ChAT_Shh_^-/-^ n=5, Nestin_Shh_^-/-^ n=4, Olig2_Shh_^+/-^ n=4, Olig2_Shh_^-/-^ n=5. Means ± SEM are shown. One-way ANOVA, Dunnett’s multiple comparison post hoc test. *p<0.05, **p<0.01, ***p<0.001. (**G**) Quantification of total numbers of MMC, HMC, and PGC MNs. Means ± SEM are shown. One-way ANOVA, Dunnett’s multiple comparison post hoc test. *p<0.05, ** p<0.01, ***p<0.001. (**H**) Ratio of each column to total thoracic MNs. Means ± SEM are shown. One-way ANOVA, Dunnett’s multiple comparison post hoc test. NS, not significant, P>0.5. (**I**) Immunostaining and quantification of late born migrating Hb9+ MNs at E12.5 brachial and thoracic segments. Shh^C/C^ n=6-7, ChAT_Shh_^-/-^ n=5, Nestin_Shh_^-/-^ n=3-4, Olig2_Shh_^+/-^ n=3-4, Olig2_Shh_^-/-^ n=5. Means ± SEM are shown. One-way ANOVA, Dunnett’s multiple comparison post hoc test. ***p<0.001. (**J**) Immunostaining and quantification of Hb9+ Caspase3+ apoptotic MNs at E12.5 brachial and thoracic segments. Shh^C/C^ n=3-6, Olig2_Shh_^+/-^ n=3, Olig2_Shh_^-/-^ n=4-5. Means ± SEM are shown. One-way ANOVA, Dunnett’s multiple comparison post hoc test. ** p<0.01, ***p<0.001. (**K**) Schematic representing Shh ablation within the genotypes and associated numbers of average MNs for brachial and thoracic sections. Scale bars, 50 μm.

Quantification of MN numbers revealed no differences in ChAT_Shh_^-/-^ and Nestin_Shh_^-/-^ compared to Shh^C/C^ controls at brachial and thoracic levels. In contrast, in Olig2_Shh_^+/-^ and Olig2_Shh_^-/-^ we find a 22% and 38%, resp. reduction in the numbers of total MNs at brachial (**Fig. 3B**), and 22% and 32% at thoracic levels compared to Shh^C/C^ controls (**Fig. 3F**). Since the production of late born MNs could be affected to a greater extent than early born MNs by the ablation of Shh from previously born VZD, we compared the relative size of the MN columns and the numbers of late born MNs just emerging from the ventricular zone at E12.5. We found that MNs attained columnar identities in normal relative proportions in all genotypes (**Fig. 3D and 3H**). However, quantification of migrating late born Hb9+ MNs showed a dose-dependent reduction of ~10% in Olig2_Shh_^+/-^ and 47% in Olig2_Shh_^-/-^ in brachial, and 23% and 47% resp. in thoracic segments, suggesting that the deficit in MN generation that we observe in Olig2_Shh_^+/-^ and Olig2_Shh_^-/-^ mice is greatest towards the end of MN production (**Fig. 3I**). We did not observe a decrease in numbers of late born migrating MNs in ChAT_Shh_^-/-^ or Nestin_Shh_^-/-^. We next examined if the earlier deficits in MN generation in Olig2_Shh_^-/-^ mice could be overcome by reduced apoptosis during the phase of programmed cell death. To this end, we found reduced levels of Caspase3+ Hb9+ brachial MNs in Olig2_Shh_^-/-^ at E12.5 indicating that reduced cell death of MNs at least in part will compensate for a reduced rate of MN production in Olig2_Shh_^-/-^ animals (**Fig. 3J**).

We associated the degree of reduction in the numbers of MNs with the tissue specific efficiency of Shh ablation in ChAT_Shh_^-/-^, Nestin_Shh_^-/-^, and Olig2_Shh_^-/-^ (**Fig. 3K**). The 80% efficient ablation of Shh from MNs at brachial levels as well as the near complete ablation of Shh from the LFP in Nestin_Shh_^-/-^, has no effect on MN numbers at E12.5. In contrast, the near complete ablation of Shh from MNs, LFP and about 50% of MFP in Olig2_Shh_^-/-^ results in a Shh dosedependent reduction of MN numbers at all levels and in all MN columns (**Fig. 3K, orange line**). Together these results reveal that MN differentiation can proceed in the absence of Shh signaling from the ventricular zone derivatives MN and LFP, but the efficacy of MN generation becomes progressively impacted by dose-dependent reductions in Shh signaling from the MFP.

### Shh from VZD sources in addition to MFP is critical for pMN domain maintenance during the onset of gliogenesis in a spinal level specific manner

Previous studies did not evaluate the contribution from individual Shh sources for the maintenance of established precursor domains after NC retraction. We therefore next investigated ventricular zone organization and precursor domain maintenance in ChAT_Shh_^-/-^, Nestin_Shh_^-/-^, and Olig2_Shh_^-/-^. We find that the relative location and distance to each other of the p3, pMN, and p0 precursor domains are preserved in ChAT_Shh_^-/-^, Nestin_Shh_^-/-^, and Olig2_Shh_^-/-^ at E12.5 (**Fig. 4A**). However, we observe a Shh source and dose-dependent, spinal level-specific decline in the numbers of Olig2+ cells in the pMN domain of ChAT_Shh_^-/-^, Olig2_Shh_^+/-^, Nestin_Shh_^-/-^, and Olig2_Shh_^-/-^ compared to Shh^C/C^ or Olig2-Cre control (**Fig. 4B and 4C**).

**Fig. 4.**
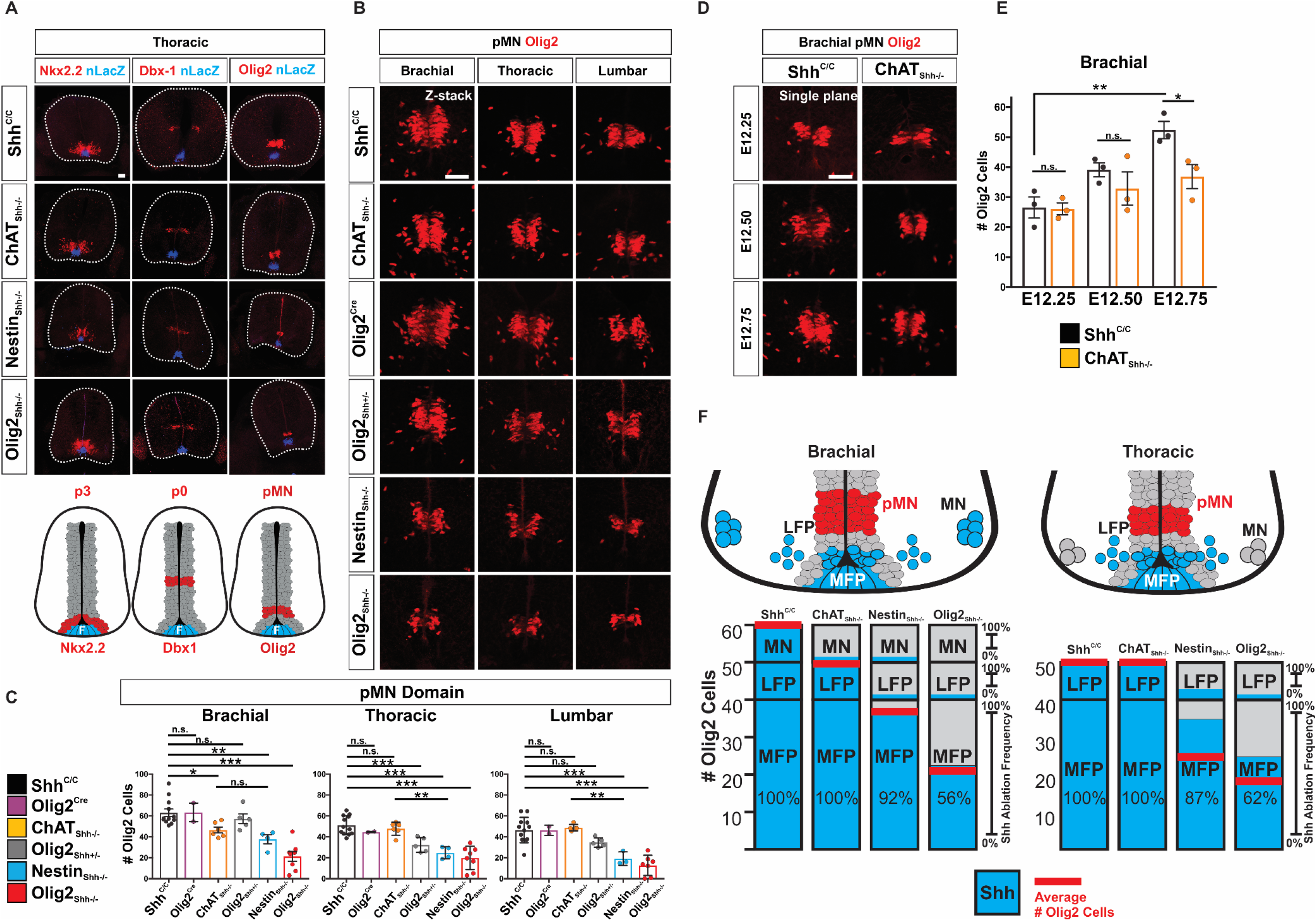
Shh from VZD in addition to MFP is critical for pMN domain maintenance in a spinal level specific manner. (**A**) Immunostaining of nLacZ with Nkx2.2, Dbx-1, and Olig2 on E12.5 thoracic sections. (**B**) Immunostaining for Olig2 in the pMN domain along the AP axis on brachial, thoracic, and lumbar sections for Shh^C/C^, ChAT_Shh_^-/-^, Nestin_Shh_^-/-^, Olig2-Cre, Olig2_Shh_^+/-^, and Olig2_Shh_^-/-^. (**C**) Quantification of numbers of Olig2 cells in the pMN domain for brachial, thoracic, and lumbar sections. Means ± SEM are shown. Shh^C/C^ n=12, Olig2-Cre n=2, ChAT_Shh_^-/-^ n=6, Olig2_Shh_^+/-^ n=5, Nestin_Shh_^-/-^ n=4, Olig2_Shh_^-/-^ n=8. One-way ANOVA, Dunnett’s or Tukey’s multiple comparison post hoc test. *p<0.05, **p<0.01, ***p<0.001. (**D**) Expansion of pMN domain at onset of gliogenesis at brachial segments between E12.25-E12.75 is reduced in ChAT_Shh_^-/-^. (**E**) Quantification of Olig2 cells in the pMN domain on brachial segments. Means ± SEM are shown. E12.25 Shh^C/C^ n=3, ChAT_Shh_^-/-^ n=3, E12.50 Shh^C/C^ n=3, ChAT_Shh_^-/-^ n=3, E12.75 Shh^C/C^ n=3, ChAT_Shh_^-/-^ n=3. Data were analyzed by Student’s t test. *p<0.05. (**F**) Schematic representing Shh ablation within the genotypes and associated numbers of average pMN Olig2 cells for brachial and thoracic sections. Blue columns indicate percent of Shh expressing cells normalized to Shh^C/C^ controls for MFP, LFP, and MNs. Grey areas indicate Cre ablation. Red bars indicate average numbers of Olig2 cells within the pMN. Scale bars, 50 μm.

In ChAT_Shh_^-/-^ we find a 26% decrease in pMN/Olig2+ cells (pMN_Olig2_^+^) at brachial levels (but not at thoracic or lumbar) consistent with expression of Shh in MNs at brachial but not yet in MNs at thoracic and lumbar levels which will begin to express Shh about one day later (**Fig. 4C**). Upon more detailed analysis, we find the effect of MN Shh to become progressively more pronounced during the rapid enlargement of the pMN that occurs at brachial segments between E12.5 to E12.75 (**Fig. 4D and 4E**). The numbers of pMN_Olig2_^+^ cells in Nestin_Shh_^-/-^ displays an anterior-posterior progressive decrease of 40% at brachial, 51% at thoracic, and 59% at lumbar segments (**Fig. 4B and 4C**). The most severely affected genotype is Olig2_Shh_^-/-^ with a decrease in pMN_Olig2_^+^ cells of 66% at brachial, 61% at thoracic, and 72% at lumbar (**Fig. 4B and 4C**).

We next associated the degree of reduction in the numbers of pMN_Olig2_^+^ cells at brachial and thoracic levels with the time of onset and tissue specific efficiency of Shh ablation in ChAT_Shh_^-/-^, Olig2_Shh_^-/-^, and Nestin_Shh_^-/-^ (**Fig. 4F**). In controls, we find about 19% more pMN_Olig2_^+^ cells at brachial than thoracic segments. The 80% efficient ablation of Shh from MNs in ChAT_Shh_^-/-^ reduces the numbers of pMN_Olig2_^+^ cells at brachial levels to those present in controls at thoracic levels while the ablation of Shh from MNs has no effect on the numbers of pMN_Olig2_^+^ cells at thoracic levels (**Fig. 4F, red line**). Nestin_Shh_^-/-^ and Olig2_Shh_^-/-^ exhibit near complete ablation of Shh from MNs and LFP, but significantly different ablation efficiencies in the MFP, resulting in a reduction of pMN_Olig2_^+^ cells at brachial levels that scales with the degree of Shh ablation from the MFP. At thoracic levels, however, we find the same magnitude in the reduction of the numbers of pMN_Olig2_^+^ cells in Nestin_Shh_^-/-^ and Olig2_Shh_^-/-^ despite of a much greater total reduction in numbers of Shh producing cells by Olig2-Cre compared to Nestin-Cre. This observation suggests that Shh produced by LFP cells, though few in number, has a disproportionate significance compared to MFP derived Shh for pMN_Olig2_^+^ cells at thoracic segments.

Next, we determined if the decreased numbers of pMN_Olig2_^+^ cells is the result of diminished initial specification or a failure of maintenance. We lineage traced Olig2 cells in Olig2_Shh_^-/-^ mutants, Olig2_Shh_^+/-^ controls, and Olig2-Cre controls using the R26mT/mG reporter allele from which myristylated GFP is expressed in all derivatives of Olig2 expressing cells and immunostained for Olig2 at E12.5 (**Fig. 5A**). We observe GFP expression in the ventral spinal cord forming a dorsal boundary at a similar relative distance to the MFP in mutants and controls but a decline in pMN_Olig2_^+^ GFP+ double positive cells in mutants compared to controls. Notably, the absence of Olig2 expressing cells within the GFP labeled area in mutants is most pronounced in the dorsal half of the pMN domain. Together, these results demonstrate that ongoing Shh signaling originating from VZD in addition to MFP are critical for the selective maintenance of pMN_Olig2_^+^ cells once the influence of notochord Shh has waned and the pMN switches to gliogenesis of OPCs.

**Fig. 5.**
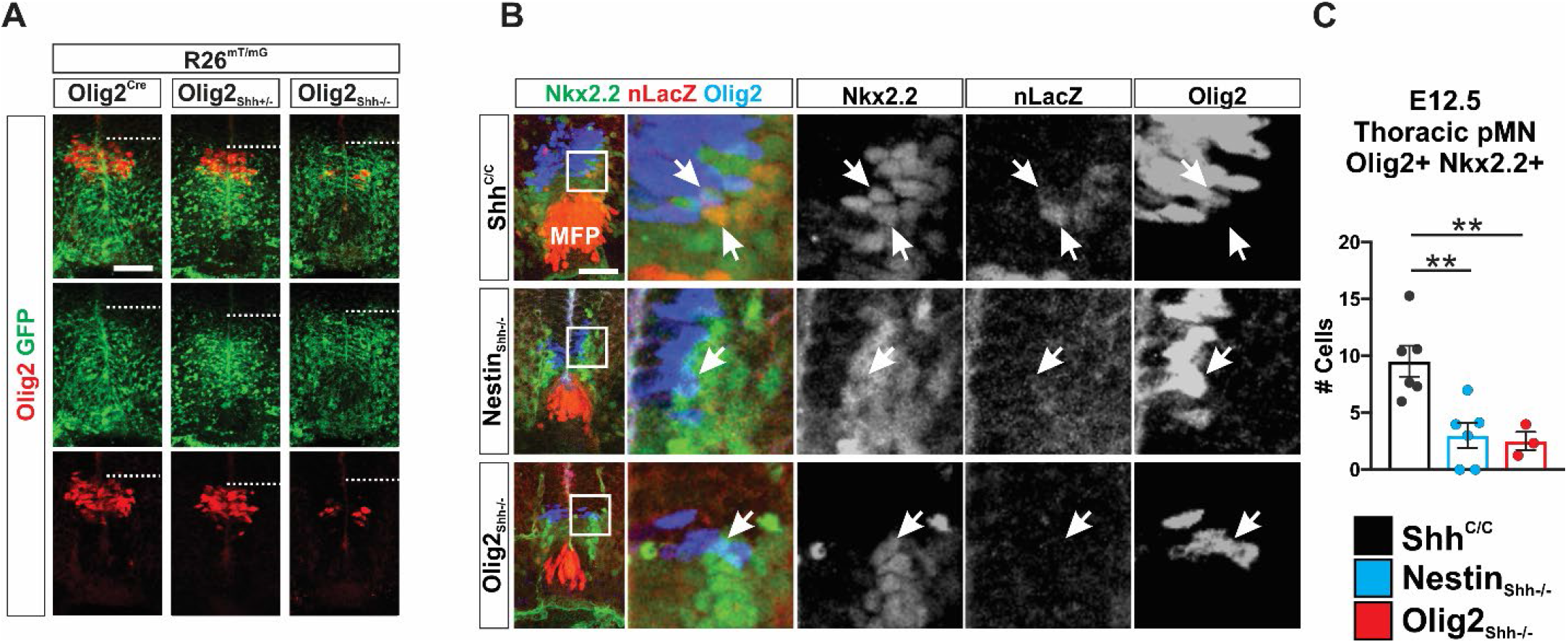
pMN_Olig2_+ expression and p* domain formation requires Shh from MFP and VZD during onset of gliogenesis. (**A**) E12.5 Lineage tracing reveals correct establishment of the pMN in Olig2_Shh_^-/-^ as indicated by dorsal boundary of R26mT/mG expression, however failure of maintenance of Olig2 in pMN as detected by immunolabeling. Scale bars, 50 μm. (**B**) p* domain is found in contact with LFP* in Shh^C/C^ controls but not in Nestin_Shh_^-/-^ and Olig2_Shh_^-/-^ as identified by immunolabeling of Olig2+ Nkx2.2 (p*) and nLacZ+ Nkx2.2+ (LFP*). (**C**) Quantification of p* domain in Shh^C/C^, Nestin_Shh_^-/-^, and Olig2_Shh_^-/-^. Means ± SEM are shown. Shh^C/C^ n=6, Nestin_Shh_^-/-^ n=6, Olig2_Shh_^-/-^ n=3. One-way ANOVA, Dunnett’s multiple comparison post hoc test. **p<0.01.

Providing further evidence for decreased pMN domain activity at the beginning of OPC production at E12.5 we find a ~3-fold reduction in the size of p* precursor domain in Olig2_Shh_^-/-^ compared to controls (**Fig. 5B and 5C**). The p* domain forms at the ventral border of the pMN domain and is marked by Olig2+/Nkx2.2+ cells (Agius et al., 2004). Interestingly, we found LFP* cells (which we define as Nkx2.2+ nLacZ+ cells) in direct contact with Nkx2.2+ Olig2+ double positive cells of the p* domain in Shh^C/C^ controls, highlighting a cyto-architectural arrangement that could underpin the disproportionate importance of LFP compared to MFPShh for the maintenance of the pMN_Olig2_^+^ cell population (**Fig. 5B**).

### Diminishment of pMN_Olig2_^+^ cells during the phase of ventral oligodendrogenesis results in reduced OPC production

Reduced Shh signaling originating from MNs, LFP, or MFP leaves the pMN domain impoverished of pMN_Olig2_^+^ cells at the beginning of oligodendrogenesis (**Fig. 4**). Nevertheless, the pMN_Olig2_^+^ cell population could recover during OPC production by increased precursor recruitment from dorsal ventricular precursor domains (Ravanelli and Appel 2015), proliferation of remaining pMN_Olig2_^+^ precursor cells, or increased differentiation and amplification of OPC fated cells that have left the pMN domain. We therefore first visualized the size of pMN_Olig2_^+^ population at the end of ventral oligodendrogenesis at E14.5. We find a moderate reduction in the numbers of pMN_Olig2_^+^ cells in ChAT_Shh_^-/-^ and an almost complete absence of pMN_Olig2_^+^ cells in Nestin_Shh_^-/-^ and Olig2_Shh_^-/-^ compared to Shh^C/C^ controls suggesting that increased recruitment of precursors to the pMN domain does not occur in mutants (**Fig. S4 and Fig. 6A**). We then determined whether pMN_Olig2_^+^ cells and/or migrating OPCs in Olig2_Shh_^-/-^ increase their rate of proliferation during the phase of OPC production. We injected EdU into pregnant dams at E11.5, E12.5, and E13.5 and quantified the numbers of pMN_Olig2_^+^ cells 24h later. We found comparable broad EdU+ labeling throughout the ventricular zone of Olig2_Shh_^-/-^ mutants and controls, suggesting overall progenitor proliferation is not affected in mutants (**Fig. 6A**). Within the pMN domain we observe a ~25% decrease in the numbers of pMN_Olig2_^+^ cells in Shh^C/C^ controls over the course of OPC production from E12.5 to E14.5 (**Fig. 6B**). In contrast, numbers of pMN_Olig2_^+^ cells in Olig2_Shh_^-/-^ mutants decline to near undetectable levels during the same period indicating a precocious exhaustion of the pMN_Olig2_^+^ cell population during OPC generation (**Fig. 6B**). We next determined the rate of proliferation of pMN_Olig2_^+^ cells in controls and mutants. In controls, we find a similar proliferative rate of about 47% of pMN_Olig2_^+^ cells at E12.5, E13.5 and E14.5. In contrast, in Olig2_Shh_^-/-^ we find the rate of proliferation to be reduced to 37% at E12.5 and E13.5, followed by a further decrease to 16% by E14.5 (**Fig. 6C**). These results indicate that the precocious exhaustion of the pMN_Olig2_^+^ cell population is associated with a reduced proliferation rate of precursor cells during OPC production which is compounded by reduced numbers of pMN_Olig2_^+^ cells that are present at the beginning of OPC production.

**Fig. 6.**
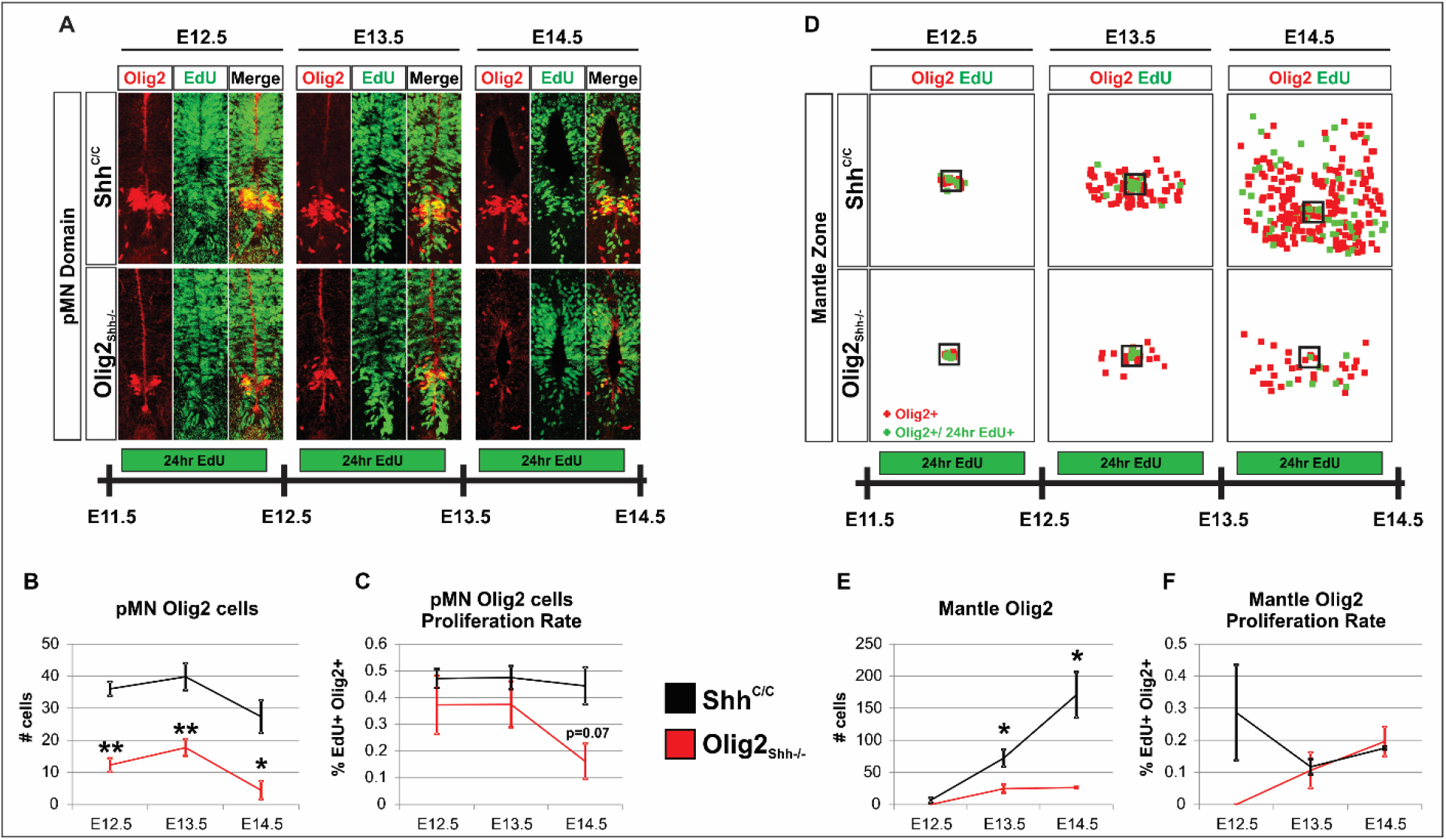
Exhaustion of pMN_Olig2+_ cells results in diminished OPC production. (**A**) pMN domain proliferation in lumbar sections labeled by EdU incorporation in 24hr intervals for E12.5-E14.5 in Shh^C/C^ and Olig2_Shh_^-/-^. (**B**) Total Olig2 cells in pMN at E12.5-E14.5. Means ± SEM are shown. Shh^C/C^ n=3-4, Olig2_Shh_^-/-^ n=3-4. Data were analyzed by Student’s t test. *p<0.05, **p<0.01. (**C**) Proliferation rate of Olig2 cells in pMN E12.5, E13.5, and E14.5. Means ± sEM are shown. Shh^C/C^ n=3-4, Olig2_Shh_^-/-^ n=3-4. Data were analyzed by Student’s t test. (**D**) Tracing of migrating Olig2 cells on representative lumbar sections with proliferation labeled by EdU incorporation in 24hr intervals between E12.5-E14.5. Box indicates pMN domain, Olig2+ cells in the mantle zone (red), Olig2+ EdU+ cells (green). (**E**) Total Olig2 cells in mantle zone at E12.5, E13.5, and E14.5. Olig2 cell numbers in Olig2_Shh_^-/-^ mantle remain reduced. Means ± SEM are shown. Shh^C/C^ n=3-4, Olig2_Shh_^-/-^ n=3-4. Data were analyzed by Student’s t test. *p<0.05. (**F**) Proliferation rate of Olig2 cells in mantle zone at E12.5, E13.5, and E14.5. Olig2 cells that have migrated out of the pMN in Olig2_Shh_^-/-^ mutants proliferate at the same rate as controls. Means ± SEM are shown. Shh^C/C^ n=3-4, Olig2_Shh_^-/-^ n=3-4.

We next tested whether OPC production in Olig2_Shh_^-/-^ recovers through increased amplification of precursors once they have emerged from the pMN. We analyzed numbers of EdU+ Olig2+ cells in the mantle zone (**Fig. 6D**). While there are very few Olig2+ cells in the mantle zone of mutants and controls at E12.5, with ongoing expansion of these cells, we find a 3-fold and 15-fold reduction of Olig2+ cells at E13.5 and E14.5, resp.in mutants compared to controls (**Fig. 6E**). The rate of proliferation among Olig2+ cells in the mantle zone in Olig2_Shh_^-/-^ is similar at E13.5 and E14.5 compared to controls suggesting that OPCs in mutants do not amplify at an increased rate compared to controls (**Fig. 6F**). The cells that do emerge from the pMN in Olig2_Shh_^-/-^ disperse as rapidly as their control counterparts, resulting in a ventral spinal cord that is populated with nascent OPCs with a 15-fold lower density compared to controls **(Fig. S5**). Additionally, we examined OPC proliferation at E14.5 in posterior thoracic and lumbar segments and found no detectable increase in proliferation rate in any of the pMN, mantle, or white matter (WM) areas (**Fig. S6**). These results reveal that the yield of the pMN domain during OPC production largely determines the numbers of OPCs that settle in white and grey matter.

Thus the scaled production of OPCs and oligodendrocyte along the anterior posterior extent of the spinal cord must at least in part be determined by the patterned expression of VZD_Shh_ that we find to be critical for the maintenance of the pMN_Olig2_^+^ precursor population.

## Discussion

Our study provides causal evidence that the sequential and patterned expression of Shh by previously specified cell types in the ventral spinal cord is critical for scaled oligodendrocyte precursor specification along the anterior posterior axis of the developing spinal cord. We took advantage of neural tube development as an established model system to reveal that Shh-expressing signaling centers become established among post mitotic derivatives of ventricular zone differentiation (VZD_Shh_) in a temporally- and spatially-patterned manner as development proceeds. We find that VZD_Shh_ signaling is critical for the production of OPCs in proportionate numbers to previously specified neurons. Our data provides genetic evidence in support of the hypothesis that extra-midline sources of Shh are critical for switching neurogenesis to gliogenesis and add a novel mechanism that contributes to pattern scaling along the anterior to posterior axis. Our observations do not rule out contributions of other well-established cell-autonomous and cell non-autonomous mechanisms that adapt morphogen function and allow pattern scaling (Houchmandzadeh, Wieschaus et al. 2005, Ben-Zvi and Barkai 2010, Hamaratoglu, de Lachapelle et al. 2011, Ben-Zvi, Fainsod et al. 2014, Uygur, Young et al. 2016), but provide a mechanism by which Shh mediated signaling from the midline in the ventral neural tube becomes progressively augmented such that Shh signaling remains a relevant instructive signal despite rapid growth.

### Sequential expression and function of VZD_Shh_

Despite its complexity, development is orchestrated by the actions of just a handful of morphogens. The temporal and spatial segregation of developmental fields that are patterned by morphogens can allow the same morphogen to play an instructive role repeatedly and at multiple anatomic regions specifying vastly different cell types and tissues. However, in the developing spinal cord all cell types are descendants of the same developmental field, the ventricular zone, yet the same morphogen, Shh, is involved in the sequential specification of multiple cell types (Dessaud, McMahon et al. 2008). The number of source tissues of Shh in the ventral spinal cord expands concomitantly with development, suggesting that “moving the Shh source” might play an important role (Danesin and Soula, 2017). We reassessed Shh expression in mice using a nuclear targeted LacZ based gene expression tracer allele and verified the principle findings in chick by RNA in situ. The advantage of the gene expression tracer allele is that the identity, location and numbers of Shh expression can be determined with single-cell resolution. Ablating Shh and nLacZ by Nestin-Cre, which is expressed in the ventricular zone, reveals that one half to one third of all Shh expressing cells in the ventral spinal cord are ventricular zone descendants at the time that Shh signaling strength within the pMN domain rises coincident with the switch from neurogenesis to gliogenesis (Dessaud, Yang et al. 2007, Balaskas, Ribeiro et al. 2012, Touahri, Escalas et al. 2012, Kicheva, Bollenbach et al. 2014, Kicheva and Briscoe 2015). While the gene expression tracer allele cannot be used to draw conclusions about the relative Shh signaling strength in the ventricular zone that is contributed by VZD_Shh_, the anatomic arrangement of these sources and the genetic ablation of Shh from these sources suggests a disproportionate effect on the activity of the pMN domain. The LFP proper extends the midline source of Shh by several cell diameters more dorsally as previously observed (Charrier et al., 2002). A subset of LFP cells, which we designate here as LFP* cells, co-express Nkx2.2 and Shh (Al Oustah, Danesin et al. 2014) and are in direct contact with cells that co-express the Shh high-threshold genes Nkx2.2+ Olig2+ and form the p* domain at the beginning of oligodendrogenesis (Fu, Qi et al. 2002, Traiffort, Zakaria et al. 2016). At the same developmental stage, we detect expression of Shh in the nascent lateral, but not medial, MN column in mice and chick. Shh can be released from the axonal as well as the dendritic compartment of neurons ((Beug, Parks et al. 2011). Hence nascent MNs could secrete Shh via dendrites or possibly trailing cellular processes close to the pMN domain. Together, the spatial and temporal pattern of VZD_Shh_ expression that we observe is consistent with the results from our selective gene ablation studies which demonstrate that these sources of Shh influence pMN domain activity, a speculation put forward previously in regard of the function of LFP_Shh_ (Danesin and Soula 2017).

A striking feature of the phenotype of Shh ablation from VZD is the temporal and qualitative segregation of the effects on MN and OPC production. VZD_Shh_ has no detectable effect on MN production or the extent of programmed cell death of MNs in our paradigms. In contrast, ablation of VZD_Shh_ results in severely reduced OPC production: (1) During the period of OPC production the initial 2-fold diminishment of pMN_Olig2_^+^ cells increases to a more than 10-fold reduction in pMN_Olig2_^+^ cells compared to controls. (2) During the same period, the numbers of Olig2+ cells that emerge from the pMN domain drop 10-fold. (3) The proliferation rate of these nascent OPCs is equal to or lower compared to their control counterparts. Together these observations reveal that the numbers of OPCs that emigrate from the pMN domain and settle in the white and grey matter are determined mainly by the size of the pMN_Olig2_^+^ precursor cell population.

The relative contribution of VZD_Shh_ to regulating the size of the pMN_Olig2_^+^ precursor cell population is spinal level specific. For example, the ablation of VZD_Shh_ results in a pMN_Olig2_^+^ loss of about 40% at brachial levels. Half of that effect at brachial levels can be attributed to the expression of MN_Shh_ since the ablation of Shh from cholinergic neurons by ChAT-Cre alone results in an almost 26% reduction in the size of the pMN_Olig2_^+^ population (**Fig. 3.2B**). The remaining size of the pMN_Olig2_^+^ population at brachial levels in the ChAT_Shh_^-/-^ spinal cord is similar to the average size of the pMN_Olig2_^+^ domain at thoracic levels in control spinal cords. Since thoracic MNs do not express Shh until the end of OPC production at E14.5, our observations reveal that the increased size of the pMN domain at brachial levels is dependent on MN_Shh_. Thus, the selective expression of MN_Shh_ at brachial and lumbar levels provides a mechanism for ensuring a proportionate increase in the production of OPCs at brachial and lumbar – compared to thoracic-levels that is matched to the increased numbers of MNs at limb levels.

### Potential mechanisms of actions of VZD_Shh_

Does Shh from different sources have distinct functions in the ventricular zone of the ventral spinal cord? Three observations support origin specific functions of Shh in the ventral spinal cord in the pMN domain: (1) MFPShh but not VZD_Shh_ influences MN production (Fig. 3). Consistent, ablation of Shh in the MFP by Olig2-Cre is complete at the beginning of MN production while ablation of Shh in VZD by Nestin-Cre only begins towards the end of MN generation (Fig. 1). In further support of a critical role of MFPShh in MN generation and consistent with previous findings (Yu, McGlynn et al. 2013), we find a Shh gene dose-dependent reduction of the numbers of MNs of similar magnitude among early and late forming MN columns as well as late born MNs still in transit at E12.5. Hence, based on the timing of Shh expression and the consistent deficit in MN production throughout the period of MN generation, the reduced rate of MN production must be associated with the reduction of midline-derived Shh rather than VZD-derived Shh. Strikingly, however, whether Shh expression is ablated from all VZDs (Nestin_Shh_^-/-^) or partially from the MFP in addition to LFP and MNs (Olig2_Shh_^-/-^), the pMN domain is left with similar strongly reduced numbers of pMN_Olig2_^+^ cells at the end of MN production (Fig. 2). Since ventral precursor domains form normally and a full complement of pMN_Olig2_^+^ cells is induced in Olig2_Shh_^-/-^ (Fig. 1), these results point to distinct functions of Shh derived from the MFP and VZDs in pMN domain activity: Our data indicates that MFPShh determines the rate of MN production while VZD_Shh_ is critical to counteract the exhaustion of the pMN_Olig2_^+^ population during MN production. (2) We find that the high threshold, Shh dependent and p* domain defining co-expression of Nkx2.2 and Olig2 (Fu, Qi et al. 2002) occurs in cells that are in close proximity to LFP* cells. Ablation of VZD_Shh_ results in a 3-fold reduction in the numbers of Nkx2.2/Olig2 expressing cells at E12.5 (Fig. 4). Additional ablation of Shh from 50% of the MFP in Olig2_Shh_^-/-^ did not increase the severity of this phenotype (Fig. 4). Thus, consistent with the anatomic juxtaposition of LFP cells to p* domain cells our gene ablation studies demonstrate that VZD_Shh_ is critical for the production of the Nkx2.2 expressing subpopulation of ventral oligodendrocyte lineage cells. (3) The dorsal most aspects of the pMN domain is almost completely devoid of Olig2+ expressing cells at the end of MN production at E12.5 in Nestin_Shh_^-/-^ and Olig2_Shh_^-/-^ (Fig. 2). Given that early pioneer OPCs elaborate cellular contacts selectively with MNs (Osterstock, Le Bras et al. 2018) it seems plausible that the LFP and MFP most distant pMN areas are served selectively by Shh from MN_Shh_. In this scenario Shh signaling from different sources might subdivide the pMN domain into sub-regions from which distinct oligodendrocyte subtypes might emerge (Dimou and Simons 2017, Ravanelli, Kearns et al. 2018) Dependent on the mode of delivery, VZD_Shh_ could also exhibit distinct modes of action: for example, LFP_Shh_ could result in maintenance of Olig2 expression in pMN precursor cells while MN_Shh_ could attract precursor cells to migrate into the pMN domain from the dorsal ventricular zone as is observed in zebrafish (Ravanelli and Appel, 2015).

Together, our data provides genetic evidence in support of the idea that morphogen signaling centers are established sequentially in the developing ventral spinal cord. These new “organizer tissues” express Shh which acts together with Shh produced by the medial floor plate to influence SVZ activity as the spinal cord grows. These Shh sources are critically involved in the switch of neurogenesis to oligodendrogenesis in the pMN domain. Endowing VZDs with morphogenic activity makes subsequent differentiation conditional to the completion of previous developmental milestones in the ventral spinal cord. Further, expression of the morphogen linked to the numbers and types of VZDs produced, provides a mechanism that could ensure scale invariance in regard of neuron and glia production along the anterior-posterior axis of the spinal cord.

## Acknowledgements

We thank Artur Kania for comments on earlier versions of the manuscript. We thank Thomas Jessell, Susan Morton, Ben Novitch, Sam Pfaff for reagents. LS performed the experiments, analyzed data and wrote the manuscript. AHK conceived the work, interpreted the results and wrote the manuscript. LS was funded in part by the Graduate Center of the City University of New York.

## Materials and Methods

### Transgenic Mice

All animal experiments were approved by the Institutional Animal Use Care Committee at CUNY. The following mouse strains were used and genotyped as described previously: Shh-nLZ^L/+^ animals (Gonzalez-Reyes et al., 2012), Chat-Cre (Rossi et al., 2011), Olig2-Cre (Dessaud 2007), Nestin-Cre (Tronche et al., 1999), Rosa26^mT/mG^ (Muzumdar et al., 2007). Mice were maintained on a C57BL/6 background. Noon on the day of the plug was considered E0.5. Mice were kept on a 12 hr dark/light cycle and the day of birth designated P1. For E12.25, E12.50, and E12.75 pMN analysis, pregnant dams were sacrificed at E12.5 according to plug date and embryos were binned into three groups (E12.25, E12.50, and E12.75), based on how many Olig2+ cells have migrated out of the pMN domain. Embryo sections which had an average of less than 5 cells migrate out of the pMN were considered 12.25, 5-20 cells migrating out – E12.50, and >20 cells migrating out – E12.75.

### In vivo EdU Assay

Pregnant dams received EdU (5-ethynyl-20-deoxyuridine, Invitrogen) dissolved in PBS by intraperitoneal injection (50 mg/kg) and sacrificed after 24 hours. Tissue sections were stained using the Click-iT Plus EdU Alexa Fluor 647 Imaging Kit (Thermofisher).

### Tissue Processing

All mice were sacrificed using an overdose of anesthetic, subjected to transcardial perfusion with 4% (w/v) paraformaldehyde (PFA) in 0.1 M PBS pH 7.4. Spinal cords and embryos were dissected, postfixed in 4% PFA for 1 hr at 4°C, cryoprotected with 30% (w/v) sucrose in 0.1M PBS for 24-48 hr, embedded and frozen in OCT medium, and stored at −80°C. Tissues were sectioned at 20 μm and collected onto glass slides.

### Immunocytochemistry and Microscopy

20um thick spinal cord cryosections were air dried for 30 min. Then sections were washed with PBS for 10 mins and with 0.3% [v/v] Triton X-100 in PBS for 20 min. Sections were then pretreated with blocking solution (10% [v/v] horse serum and 0.3% [v/v] Triton X-100 in PBS) for 90 mins and incubated with primary antibodies overnight at 4C. The next day, following 3 PBS washes the sections were incubated with secondary antibodies for 2 hr at room temperature. A list of all antibodies and compounds used is provided in a table. For cell counts, at least three sections per animal from at least three mice were examined, unless otherwise noted. Images were acquired using a Zeiss LSM880 confocal microscope.

Statistical analysis was performed using Prism 7 (Graphpad Software Inc.) Analysis of multiple groups was made using one-way ANOVA followed by the Tukey or Dunnett’s post hoc analysis tests. For 2-groups analyses, unpaired Student’s t test was used. The data are presented graphically as: *(p < 0.05), **(p < 0.01), and ***(p < 0.001).

| REAGENT | SOURCE | IDENTIFIER |
| --- | --- | --- |
| **Antibodies** |  |  |
| Chicken polyclonal anti beta Galactosidase | Abcam | Cat# ab9361,<br>RRID:AB_307210 |
| Goat polyclonal anti beta Galactosidase | Biogenesis | Cat# 4600-1409,<br>RRID:AB_2314510 |
| Goat polyclonal anti Choline acetyltransferase | Millipore | Cat# AB144P,<br>RRID:AB_2079751 |
| Rabbit polyclonal anti Olig2 | Millipore | Cat#AB9610,<br>RRID:AB_570666 |
| Mouse monoclonal anti Nkx2.2 | DSHB | Cat# 74.5A5,<br>RRID:AB_531794 |
| Rabbit polyclonal anti nNos | Immunostar | Cat# 24431,<br>RRID:AB_572255 |
| Mouse monoclonal Ankyrin G (AnkG) (463) | Santa Cruz | Cat# sc-12719,<br>RRID:AB_626674 |
| Mouse monoclonal Lim3 (Lhx3) | DSHB | Cat# 67.4E12,<br>RRID:AB_2135805 |
| Rabbit polyclonal Lim1 | Gift Jessell Lab | N/A |
| Guinea Pig polyclonal Hb9 | Gift Jessell Lab | N/A |
| Rabbit polyclonal FoxA2 | Gift Jessell Lab | N/A |
| Rabbit polyclonal Pax6 | Gift Jessell Lab | N/A |
| Rabbit polyclonal Dbx1 | Gift Jessell Lab | N/A |
| Rabbit polyclonal Isl1/1 | Gift Jessell Lab | N/A |
| Rabbit polyclonal anti Nkx2.2 | Gift Jessell Lab | N/A |
| Donkey anti rabbit Alexa 488 | Jackson ImmunoResearch | Cat# 711-545-152, RRID:AB_2313584 |
| Donkey anti mouse Alexa 488 | Jackson ImmunoResearch | Cat# 715-545-150, RRID:AB_2340846 |
| Donkey anti guinea pig Alexa 488 | Jackson ImmunoResearch | Cat# 706-545-148, RRID:AB_2340472 |
| Donkey anti goat Alexa 488 | Jackson ImmunoResearch | Cat# 705-545-147, RRID:AB_2336933 |
| Donkey anti rabbit Cy3 | Jackson ImmunoResearch | Cat# 711-165-152, RRID:AB_2307443 |
| Donkey anti mouse Cy3 | Jackson ImmunoResearch | Cat# 715-165-150, RRID:AB_2340813 |
| Donkey anti guinea pig Cy3 | Jackson ImmunoResearch | Cat# 706-165-148, RRID:AB_2340460 |
| Donkey anti rat Cy3 | Jackson ImmunoResearch | Cat# 712-165-153, RRID:AB_2340667 |
| Donkey anti rabbit Alexa 594 | Jackson ImmunoResearch | Cat# 711-585-152, RRID:AB_2340621 |
| Donkey anti mouse Alexa 594 | Jackson ImmunoResearch | Cat# 715-585-150, RRID:AB_2340854 |
| Donkey anti guinea pig Alexa 594 | Jackson ImmunoResearch | Cat# 706-585-148, RRID:AB_2340474 |
| Donkey anti rat Alexa 594 | Jackson ImmunoResearch | Cat# 712-585-153, RRID:AB_2340689 |
| Donkey anti chicken Cy5 | Jackson ImmunoResearch | Cat# 703-175-155, RRID:AB_2340365 |
| Donkey anti goat Cy5 | Jackson ImmunoResearch | Cat# 705-175-147, RRID:AB_2340415 |
| Donkey anti mouse Cy5 | Jackson ImmunoResearch | Cat# 715-175-150, RRID:AB_2340819 |
| Donkey anti guinea pig Alexa Cy5 | Jackson ImmunoResearch | Cat# 706-175-148, RRID:AB_2340462 |
| **Chemicals** |  |  |
| X-Gal | Roche | 10745740001; CAS 7240-90-6 |
| DAPI | Sigma Aldrich | CAS 28718-90-3 |
| **Commercial Assays** |  |  |
| Click-iT EdU | Thermofisher | C10340 |
| **Mouse Strains** |  |  |
| Mouse: Shh<sup>tm1Ahk</sup> | Gonzalez-Reyes L, et al., 2012 | MGI:5440762 |
| Mouse: B6;129S6-Chat<sup>tm1(cre)Lowl/J</sup> | The Jackson Laboratory | JAX: 006410 |
| Mouse: B6.Cg-Tg(Nes-cre)1Kln/J | The Jackson Laboratory | JAX: 003771 |
| Mouse: Olig2<sup>tm1(cre)Tmj</sup> | Dessaud E, et al., 2007 | MGI:3774124 |
| Mouse: Gt(ROSA)26Sor<sup>tm4(ACTB-tdTomato,-EGFP)Luo/J</sup> | The Jackson Laboratory | JAX: 007576 |

---

**Fig. S1.**
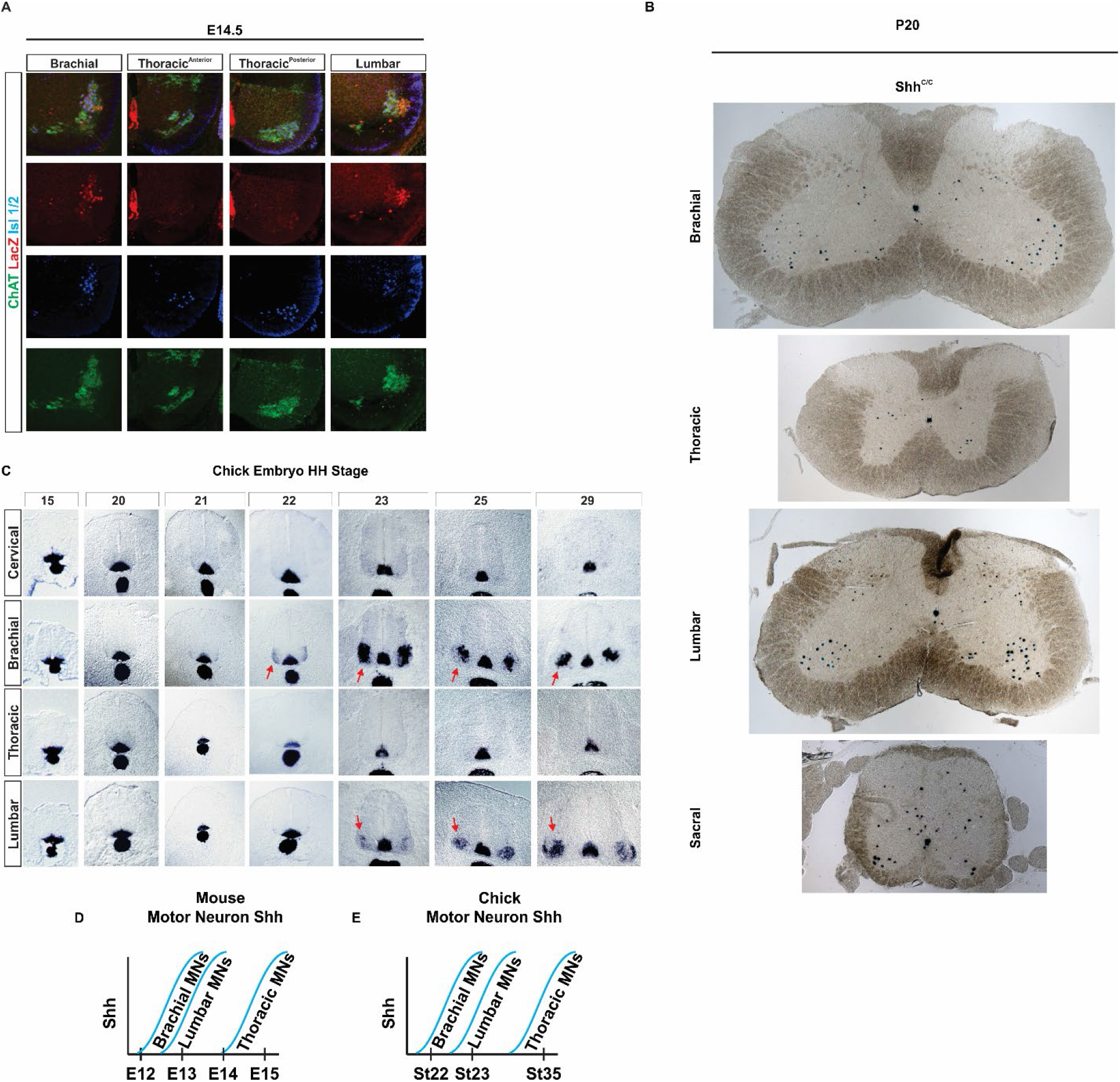
Shh expression in MNs. (**A**) Shh expression by MNs along AP axis at E14.5. Immunostaining colocalization of nLacZ with MN markers Isl1/2 and ChAT. Revealing that thoracic MNs express Shh much later than brachial and lumbar. (**B**) X-gal staining revealing Shh expression pattern throughout AP axis of P20 control Shh^C/C^ spinal cords at brachial, thoracic, lumbar, and sacral segments. (**C**) In situ hybridization for Shh in developing chick neural tube demonstrating comparable timing and pattern of Shh expression by MNs as in mouse. (**D and E**) Timeline of Shh expression by MNs in (**D**) mouse and (**E**) chick.

**Fig. S2.**
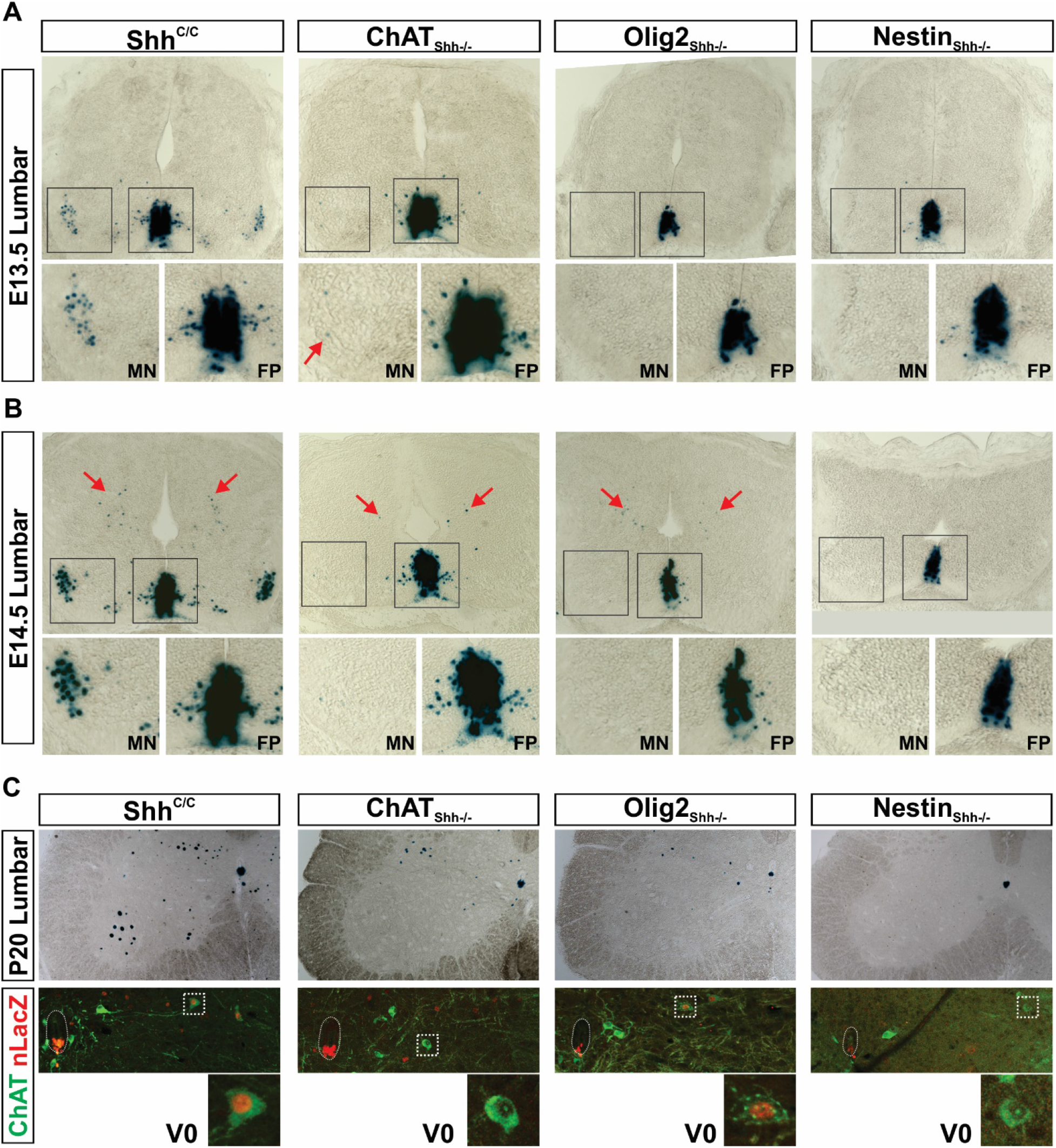
Ablation of Shh from MFP and VZD sources. (**A and B**) X-gal staining revealing ablation of Shh at lumbar spinal cord at (**A**) E13.5, and (**B**) E14.5. Arrows in B point to a dorsal Shh source located near the ventricular zone appearing at E14.5. (**C**) X-gal staining of P20 spinal cords revealing loss of Shh expression from MNs and V0 cholinergic interneurons in ChAT_Shh_^-/-^, MNs and descendants of neurons ventral to the Olig2 domain in Olig2_Shh_^-/-^, and all neuronal sources in Nestin_Shh_^-/-^.

**Fig. S3.**
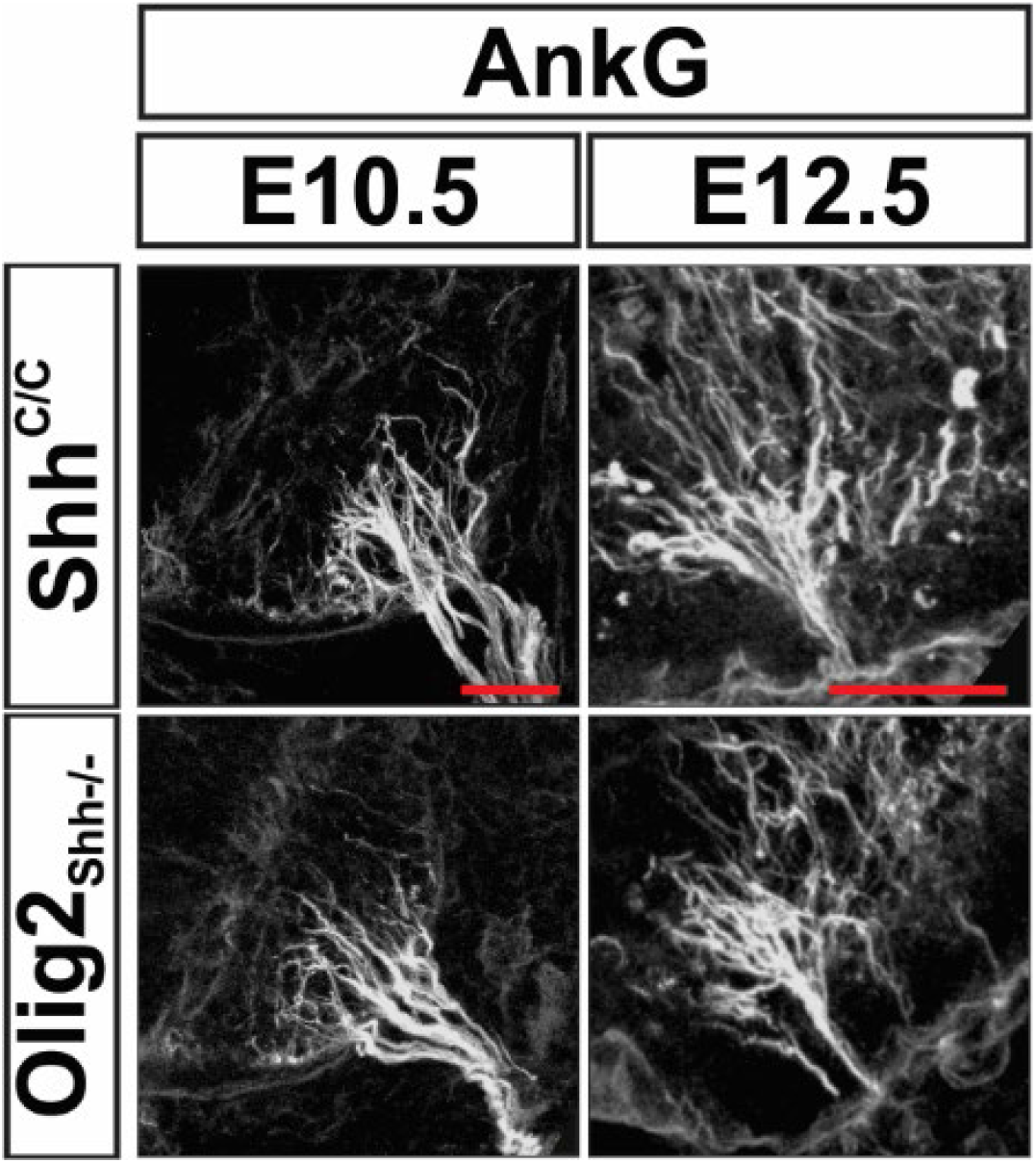
AnkG staining at E10.5 and E12.5 revealing correct MN axon fasciculation and exit from ventral horns in Olig2_Shh_^-/-^

**Fig. S4.**
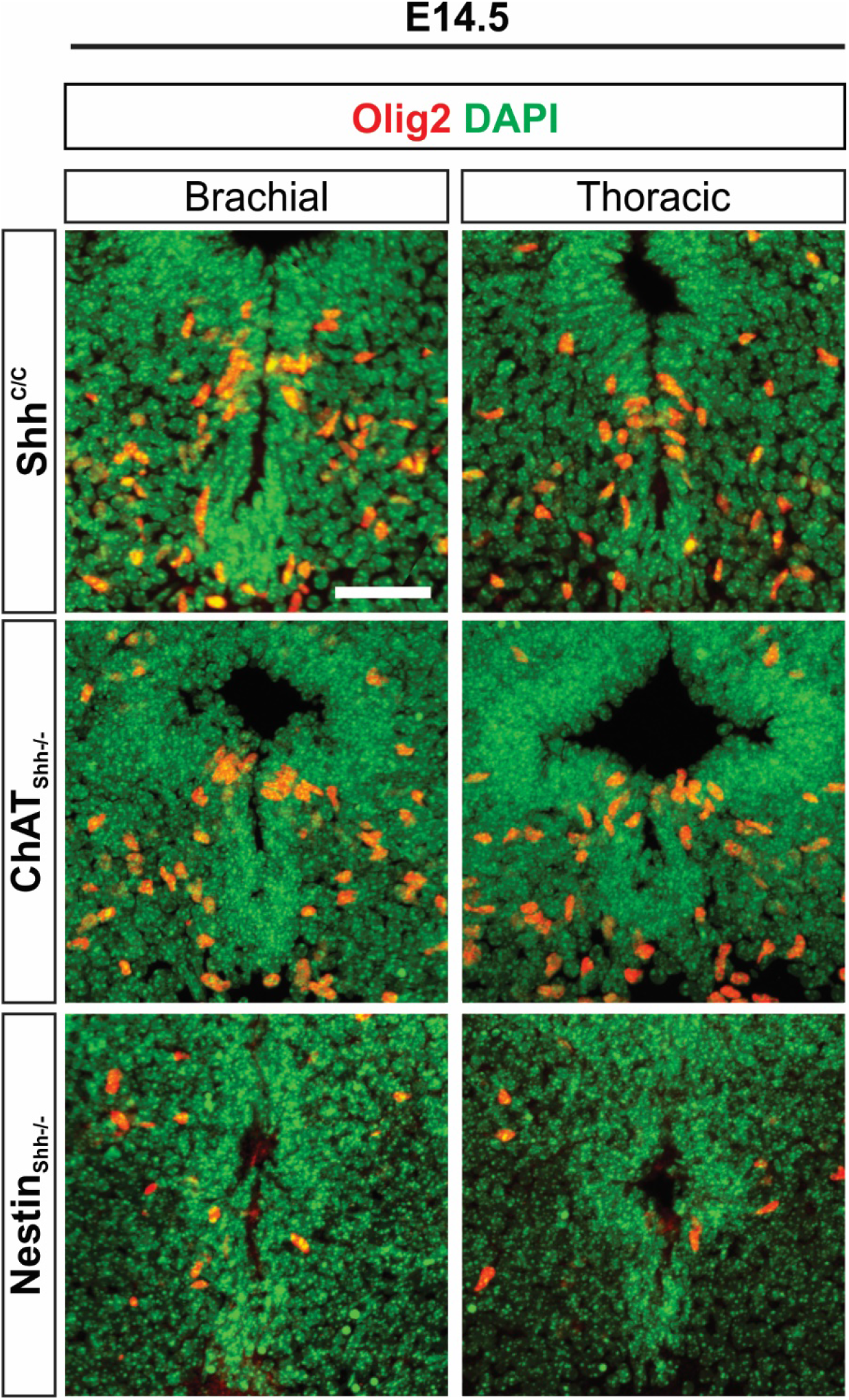
Immunostaining of Olig2 and DAPI on brachial and thoracic segments reveals the continued depletion of Olig2 cells from the pMN of Nestin_Shh_^-/-^ but not ChAT_Shh_^-/-^ at E14.5.

**Fig. S5.**
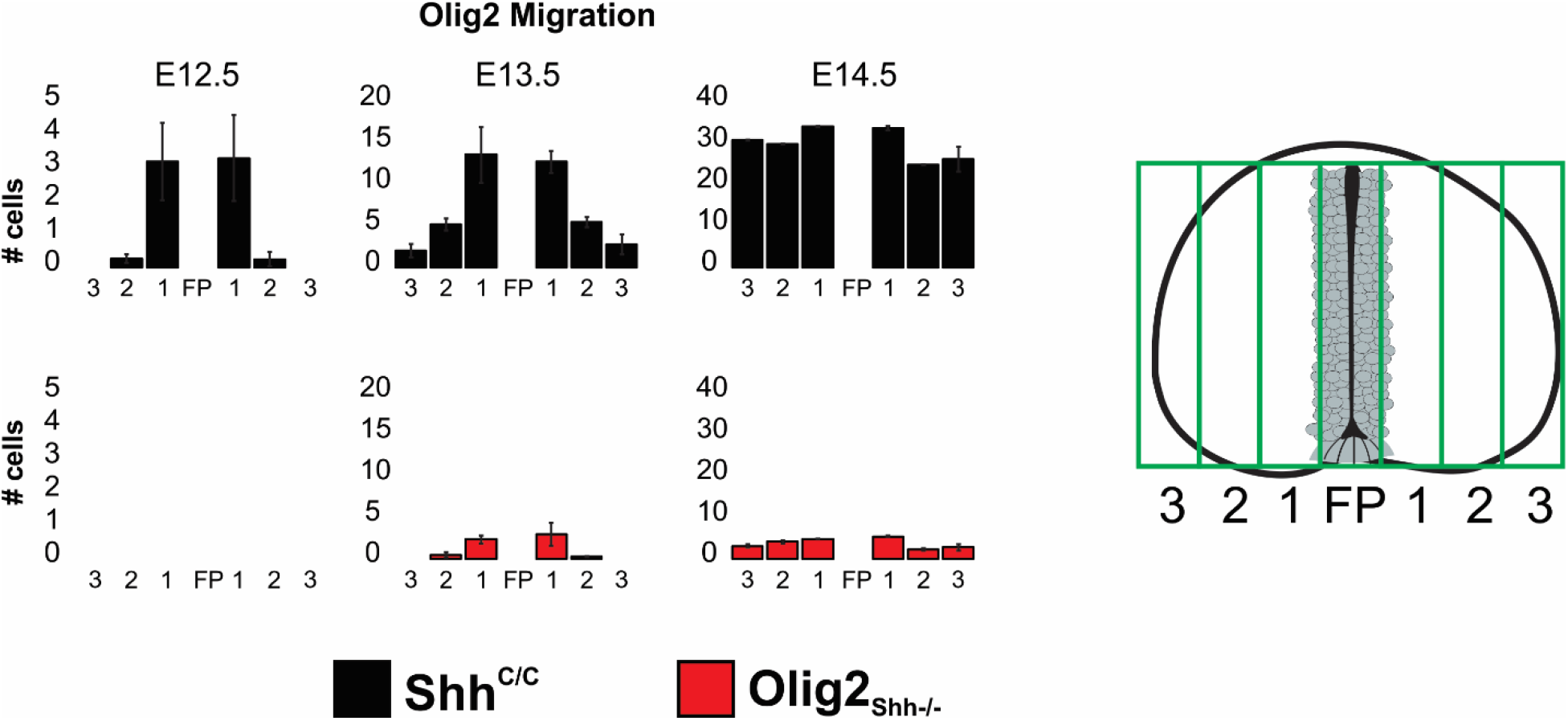
Analysis of migrating Olig2 cell dispersion in Olig2_Shh_^-/-^ mutant and control embryos. Lumbar spinal cord sections from E12.5-E14.5 were binned into 6 zones excluding the ventricular zone, and numbers of Olig2 cells in each zone were quantified. Means ± sEM are shown. Shh^C/C^ (n = 3-4 embryos), Olig2_Shh_^-/-^ (n = 3 embryos).

**Fig. S6.**
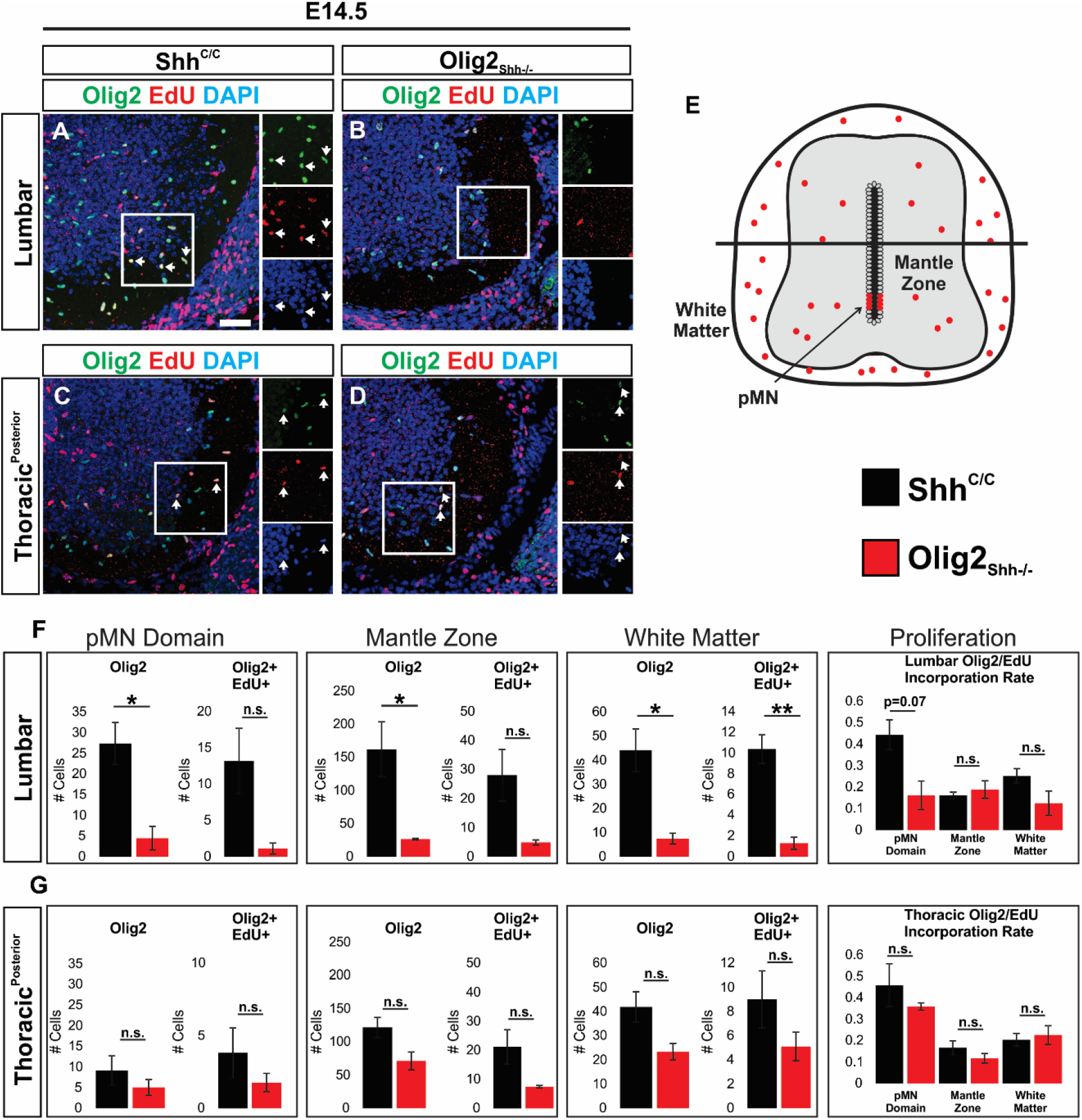
OPCs are reduced and do not increase proliferation rate. (**A-D**) Immunostaining on E14.5 sections for Olig2, EdU, and DAPI at (**A and B**) lumbar and (**C and D**) posterior thoracic. Arrows indicate co-expression of Olig2 and EdU. (**E**) Schematic depicting areas analyzed. (**F and G**) 24hr pulse chase with a single EdU injection to label proliferating Olig2 cells. Total Olig2 cells and Olig2+ Edu+ double positive cells are reduced in the pMN, mantle zone, and white matter at both (**F**) lumbar and (**G**) posterior thoracic segments in Olig2_Shh_^-/-^ compared to control. Shh^C/C^ (n = 3 embryos), Olig2_Shh_^-/-^ (n = 3 embryos). Means ± SEM are shown. Data were analyzed by Student’s t test. *p < 0.05, **p<0.01. Scale bars, 50 μm.

## Notes

### Competing Interest Statement

The authors have declared no competing interest.

